# The enhancement of electricity generation using cellulose based on ternary microbial consortium

**DOI:** 10.1101/2024.04.26.591402

**Authors:** Shengchao Gao, Tingfang Mai, Yi Ding, Zhiwei Wang, Xinxin Fan, Yue Zhang, Gongwei Liu, Ying Liu

**Author notes:** Corresponding author. Current address: Shaanxi Key Laboratory of Agricultural and Environmental Microbiology, College of Life Sciences, Northwest A&F University, No. 22 Xinong Road, Yangling, Shaanxi Province, P.R. China, 712100,. (Y. Liu). College of Animal Science and Technology, Northwest A&F University, No. 22 Xinong Road, Yangling, Shaanxi Province, PR China 712100. (GW. Liu).

## Abstract

Cooperation between microorganisms is crucial to design an efficient inoculum for enhancing the electricity-producing ability of carboxymethyl cellulose (CMC) fed bioreactors. In the present study, the influence of microbial mutualistic interactions and electricity generation capability were investigated by designing several co-culture and ternary culture systems. It was found that a ternary culture system of *Cellulomonas* Lsc-8, *Bacillus subtilis* C9 and *Geobacter sulfurreducens* PCA was used to efficiently convert cellulose into electricity. The maximum current density of 796 ± 30 µA**·**cm^-2^ were achieved by the ternary culture, which were much higher than that *Geobacter sulfurreducens* PCA using acetate and co-culture systems to utilize CMC in bioreactors, respectively. In this consortium, *Cellulomonas* Lsc-8, and *Bacillus subtilis* C9 simultaneously digested CMC to produce acetate and secreted riboflavin as an electron shuttle; *Geobacter sulfurreducens* PCA utilized acetate to generate electricity. The introduction of *Bacillus subtilis* C9 further promoted the degradation of CMC and secreted more riboflavin to enhance electricity generation of the ternary culture. This work suggested that the synergistic interaction between interspecies in microbial consortia is emergent in designing specific community for achieving maximum power generation using CMC as substrate. This research shows new insight into the design of more efficient, stable, and robust microbial consortia applicable in waste treatment and power generation.

**IMPORTANCE:** Microbial fuel cells (MFCs) may benefit from microbial consortia that efficiently convert carbon sources to electricity. A key challenge with this system is how to manage microbial community assembly to maximize electricity generation. Herein, we constructed and tested a three-species microbial consortium to enhance conversion of cellulose to electricity. *Cellulomonas* Lsc-8 and *Bacillus subtilis* C9 efficiently converted cellulose to acetate (electron donor) and riboflavin (electron shuttle), which enabled *Geobacter sulfurreducens* to generate electricity. This study laid the foundation for design of more efficient, stable, and robust microbial consortia for waste treatment and energy applications.

---

Microbial fuel cells (MFCs) are eco-friendly and emerging technology that use electrochemical active biofilm (EAB) as biocatalysts to oxidize organic compounds and product sustainable bioelectricity (1–4). A wide variety of substrates including single substrate and complicated compounds have been tested as anolyte in MFCs for electricity generation, such as municipal waste liquor, industrial wastewaters, and starch(5–8). Besides, lignocellulosic biomass has attracted great attention for use in MFCs due to its the most abundant and biodegradable biomass on earth (9, 10). Microbial fuel cells use microorganisms as biocatalysts to convert the chemical energy stored in lignocellulose into electricity (11). Therefore, designing cellulose-fed MFCs with highly efficient electricity-producing capability has great significance for current generation (12).

Recently, several studies have used undefined mixed-cultures as anode bio-catalyst in MFCs for electricity generation from cellulose (13–16). Ishii et al. (13) have successfully enriched a highly efficient mixed culture to generate electricity from rice paddy soil with a maximum current of 0.3 mA. Gregoire et al. (15) have explored the potential for converting lignocellulosic biomass directly to electricity in a MFCs with a maximum power density of 230 mW/m^3^. Yoshimura et al. (16) have found that mixing rice bran with pond sediment can increase the total charge of MFCs.

However, in aforementioned research, undefined mixed cultures could not generate significant current and it is difficult to be further optimized owing to the complex metabolites and interaction mechanisms. Several studies have focused on single bacteria strain using cellulose as sole carbon source to produce electricity, which include *Enterobacter cloacae* , *Cellulomonas fimi*, *Cellulomonas* strain Lsc-8 (17–19). However, such difunctional microbes are not efficiently utilized complex substrates (20, 21) and achieved very low electricity generation.

Synthetic microbial consortia with a defined composition and controllable functions hold great promises to solve these issues (22), there have been a few reports on defined co-culture for power generation (23–26). The catabolism process of complex natural energy sources by designing two-species consortia (*Escherichia coli* and *Shewanella oneidensis*) can enhance the magnitude of bioenergy generation(23). Aiyer (25) have used a defined co-culture to improve power generation in MFCs with a maximum power density of 190.44 mW m^-2^. Ren et al. (26) have analyzed the electricity generation of cellulose-fed MFCs and the maximum power density was 0.0153 mW**·**cm^-2^. Cao et al. (27) have designed two-species consortia including strain Lsc-8 and *Geobacter sulfurreducens* PCA as a biocatalyst for power generation with CMC, the maximum current density achieved by the co-culture with CMC was 559 μA cm^−2^. In order to enhance the electricity generation of MFC using cellulose, the defined efficient microbial consortia is needed.

We herein designed a three-species culture including *Cellulomonas* Lsc-8*, Bacillus subtilis* C9 *and Geobacter sulfurreducens* PCA for increasing electricity generation using cellulose. The chosen three species were correspondingly based on electrogenic properties and ability to utilize cellulose (19, 28, 29). *Cellulomonas* Lsc-8 and *Bacillus subtilis* C9 had the ability to convert cellulose into acetic acid, and the produced acetate was utilized by *G. sulfurreducens* PCA. Compared with the different co-culture systems (Lsc-8 and *G. sulfurreducens* PCA, *Bacillus subtilis* C9 and *G. sulfurreducens* PCA, Lsc-8 and *Bacillus subtilis* C9), The maximum current density achieved by the ternary culture with CMC was 796 ± 30 µA·cm^-2^, which were increased to 1.57, 1.43, and 22.36 times to compared with those co-culture systems using CMC, respectively. Meanwhile, the key metabolite products, including acetate and riboflavin, were quantified by high performance liquid chromatography (HPLC) to explore the biodegradation process of cellulose in the different co-culture systems and ternary system during electricity generation. Besides, from fluorescence detection, it indicated that *Bacillus subtilis* C9 can secrete riboflavin as an electron transport mediator. This research shows that the high electricity generation ability using the ternary culture resulted from the “division-of-labor” based cooperation to efficiently convert cellulose and the promoted extracellular electron transfer pathway in the ternary culture after introducing *Bacillus subtilis* C9.

## RESULTS AND DISCUSSION

### The electricity generation performance of the defined co-cultures and ternary system

Strain Lsc-8 could produce a current density of 5.10 μA cm^−2^ from CMC in the three-electrode system (Fig. S1 A), which is consistent with the previous reported on *Cellulomonas* Lsc-8 (Fig. S2 A) (27). But the electricity production was very low. Therefore, A co-culture system (strain Lsc-8 and *G. sulfurreducens* PCA) was defined to improve the electricity production. The current density was 506 ± 10 μA cm^−2^ (Fig. 1A curve c), which was similar to that obtained with *G. sulfurreducens* PCA and acetate in a bioreactor (Fig. 1A curve e). However, *G. sulfurreducens* PCA could not use CMC to generate electricity (Fig. 1A curve d). These results indicated that the co-culture system could utilize CMC to generate electricity and that the maximum current density (506 ± 10 μA cm^−2^) of the co-culture system was far higher than that of strain Lsc-8 (5.10 µA cm^−2^).

**Fig. 1.**
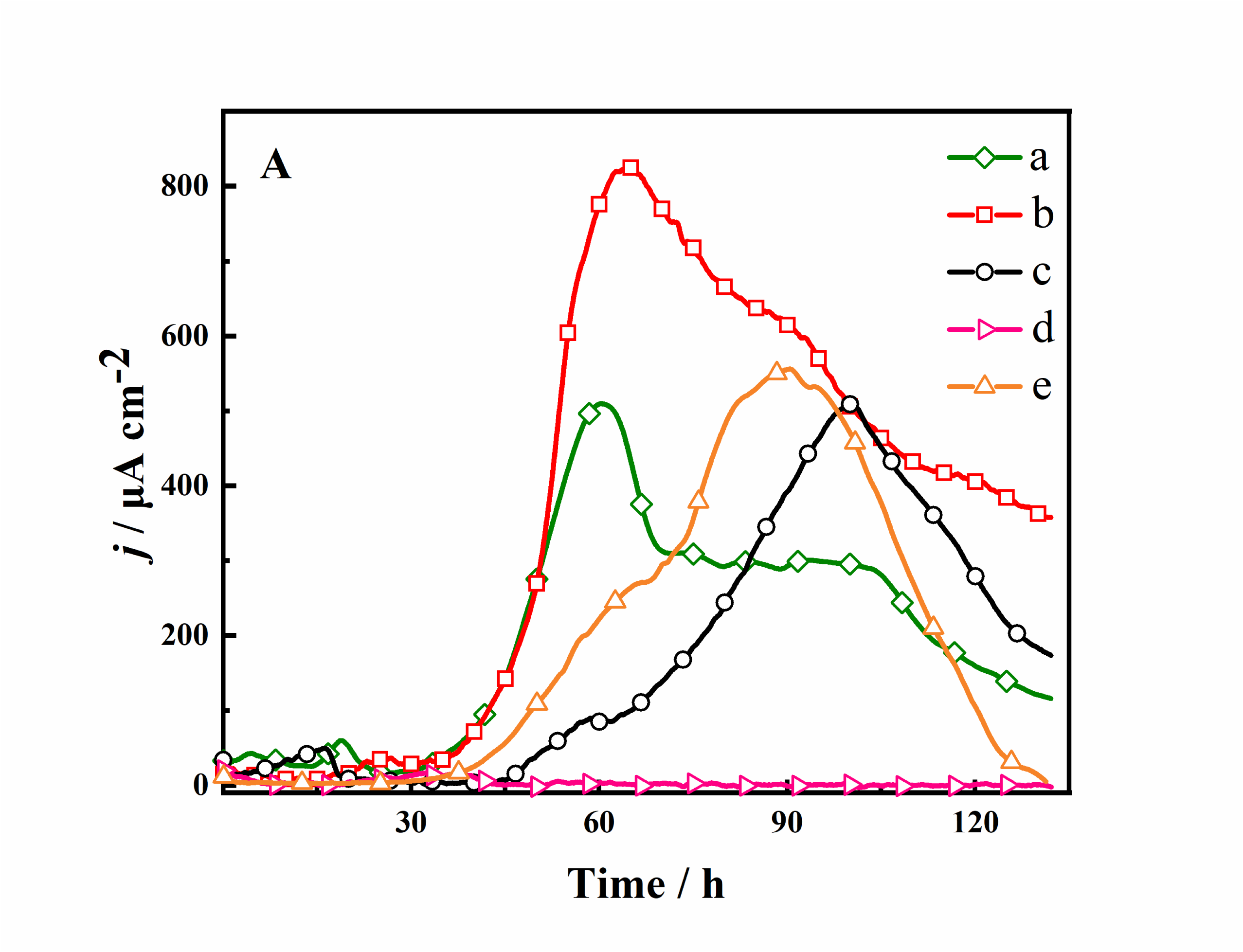
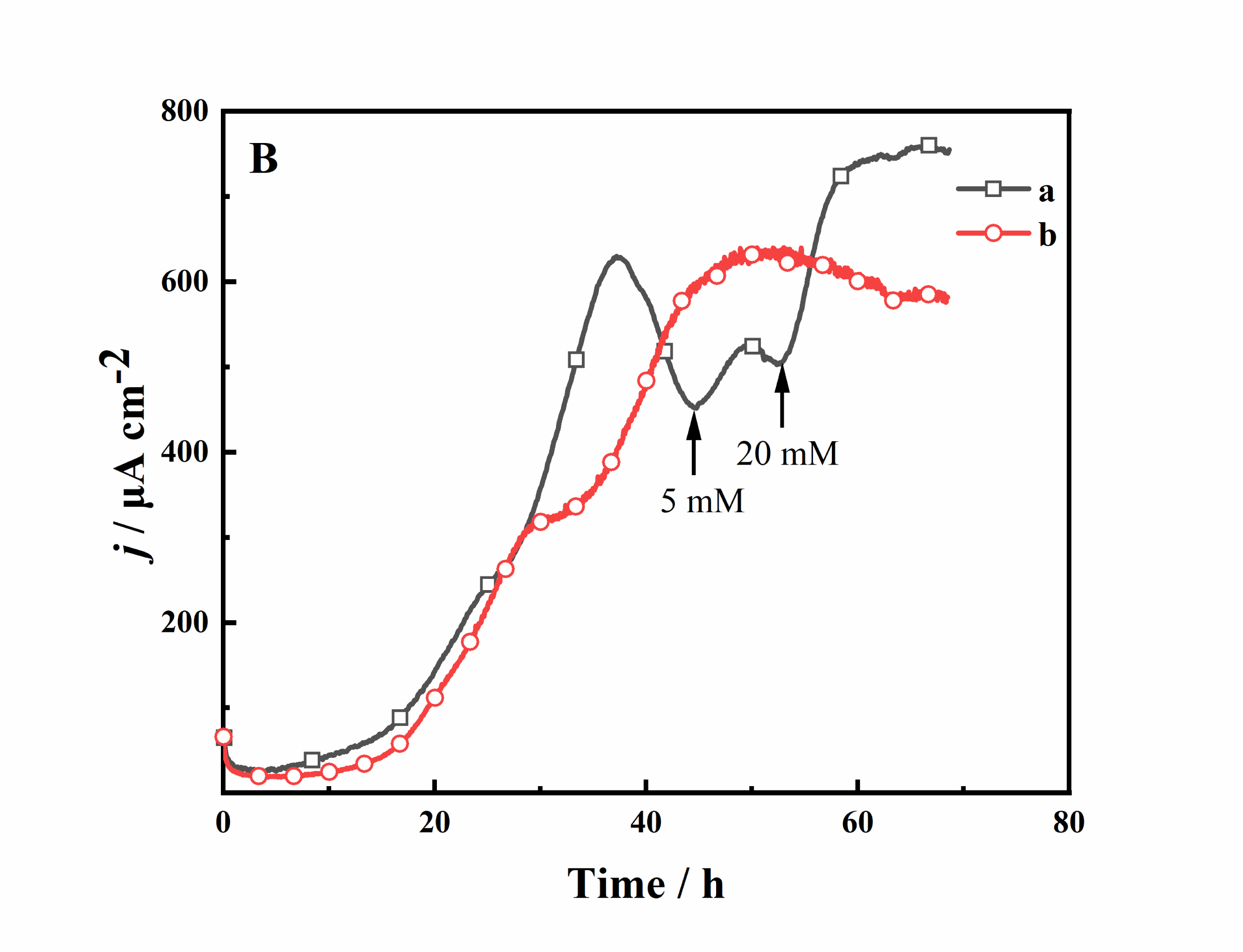
(A) The current generation of bioreactors inoculated with different bacteria with CMC or acetate as carbon source. (curve a, chronoamperometric curve of E-BG using CMC; curve b, chronoamperometric curve of E-BLG using CMC; curve c, chronoamperometric curve of E-LG using CMC; curve d, pure culture of *G. sulfurreducens* PCA with CMC; curve e. *G. sulfurreducens* PCA with acetate). (B) Chronoamperometric curves of bioreactor inoculated with the co-culture with CMC (curve a) and inoculated with *G. sulfurreducens* PCA with acetate (curve b) when the initial addition of riboflavin. The arrow indicates the addition of acetate in the bioreactor.

In order to investigate the reasons for the limitation of the electrical production performance of the co-culture system. Riboflavin and acetate (5 and 20 mM) were added to the co-culture system to investigate the impact on the bioelectricity generation, respectively. When 50 nM riboflavin was initially added to the bioreactor, the maximum current density of the co-culture system with CMC was 630.90 μA cm^−2^ (Fig 1B curve a), which was higher than that the maximum current density of bioreactor without adding riboflavin (Fig. 1A curve c). Besides, When the current density was reduced, the different concentrations of acetate (5 mM and 20 mM) were added in the bioreactor, respectively. It is shown from curve a in Fig. 1B that the current density recovered to 524.28 μA cm^−2^ and 759.25 µA cm^−2^, respectively. Besides, the maximum current density of *G. sulfurreducens* PCA with acetate was 633.25 μA cm^−2^ when 50 nM riboflavin was initially added to the bioreactor (Fig 1B curve b), which was higher than that the maximum current density of bioreactor without adding riboflavin (Fig. 1A curve e). These results indicated that riboflavin and substrate were the limiting factors to further improve the power production performance of the co-culture system.

It has been reported that *Bacillus* spp. has the effectively lignocellulose degradation ability and the production of riboflavin (30, 31). Therefore, *B. subtilis* was introduced to improve the degradation of CMC and enhance productivity. First, the cellulase activity and residual cellulose of the single and co-cultures were analyzed by DNS assay and the phenol-sulfuric acid method.

The linear relationship (R^2^= 0.9976) between glucose concentrations and solution absorbances at 540 nm with glucose concentrations of 0 - 1 mg·mL^-1^ were shown in Supporting Information Fig. S3. The co-cultures showed the highest cellulase activity (1.10 ± 0.0075 U·mL^-1^) followed by *B. subtilis* C9 (0.86 ± 0.0058 U mL^−1^), which were 5.5 times than that of strain Lsc-8 (Table S4). *B. subtilis* C9, the most competitive cellulase producer, was introduced to improve the degradation of CMC and enhance power generation. Out of our expectation, the co-culture system including strain Lsc-8 and *B. subtilis* C9 could use CMC to produce a current density of 35.60 ± 2.86 μA cm^−2^ by introducing *B. subtilis* C9 (Fig. S1 B), which was increased to 6.98 times to compared with the maximum current density of *Cellulomonas* Lsc-8. Therefore, *B. subtilis* C9 might enhanced the degradation of cellulose and could secrete electron shuttles to promote electron transfer between bacteria.

Furthermore, the various co-cultures and ternary culture were inoculated in bioreactors using CMC as electron donor, respectively. It was shown from Fig. 1A that the bioreactor inoculated with different inocula exhibited different peak current density. The maximum current density of E- BLG, E-BG, and E-LG using CMC was 796 ± 30 µA cm^-2^ (Fig. 1A curve b), 556 ± 40 µA cm^-2^ (Fig. 1A curve a), and 506 ± 10 µA cm^-2^ (Fig. 1A curve c), respectively. On the other hand, *G. sulfurreducens* PCA produced the maximum current densities of 539 ± 15 µA cm^-2^ when fed with acetate (Fig. 1 A curve e) but could not use CMC to generate electricity (Fig. 1 A curve d). It can be seen that the maximum current density achieved by E-BLG with CMC were increased to 1.43,1.57, 22.36, and 1.47 times to compared with those co-culture systems (E-BG, E-LG, and E- LB) using CMC and *G. sulfurreducens* PCA using acetate, respectively. These results implied that the various co-cultures and ternary culture systems could utilize CMC to generate electricity and that the maximum current density (796 ± 30 μA cm^−2^) of the ternary culture using CMC was much higher than those other cultures. The ternary consortium also achieved the highest CMC– bioelectricity generation comparing with previously reported results for the co-culture system (Table S3) (32–35).

### Evidence for Electron shuttles Involved in E-BG

Electron shuttles can act as electron carrier to improve the rate of electron transfer from bacteria and bacterial surfaces to conductive materials (36). It has been reported that strain Lsc-8 can secrete riboflavin to improve the electron transfer (19). In order to exclude the influence of *Cellulomonas* Lsc-8. Meanwhile, the maximum current density of the E-BG using CMC was similar than that of *G. sulfurreducens* PCA using acetate. Therefore, we investigated whether there was electron mediator in E-BG. The CV and DPV were studied to revel the electron shuttles for E-BG. When the biofilm of E-BG growth at the maximum current density using chronoamperometry, the medium with different components was changed and the CV at scan rate of 5 mV s^-1^ were recorded. As shown in curves a-d Fig. S3, all voltammograms showed typical sigmoidal shape. The catalytic current was 537.52 μA cm^−2^ when the E-BG reached the maximum current density (curve a Fig. S3). Furthermore, after addition of fresh medium containing CMC to replace the original medium caused an immediately decline in catalytic current to 58.7% (curve b Fig. S3), which may be due to the lack of electron mediator and planktonic cells in the medium. It has been reported that removal of riboflavin-containing medium reduced the maximum current of oxidation process by *Shewanella oneidensis* MR-1 (37). However, the catalytic current was restored to that of its original level when the original medium was added back to the bioreactor (curve c Fig. S3). In order to remove the effects of planktonic bacteria, the original medium was centrifuged and returned to bioreactor, the catalytic current decreased to 23% of its original level (curve d Fig. S3), which may be due to the removed planktonic cells adsorb a percentage of the electron mediator. We will study it in detail in later work. These results indicated that the medium contained compounds that act as electron shuttle to enhance the rate of electron transfer.

To further investigate whether the electron shuttles involved in E-BG was related to strain *Bacillus subtilis* C9, DPV was adopted to further confirm the electron shuttles contained in medium with *Bacillus subtilis* C9 in running 100 h to 172h (Fig. 2A). It was shown that when scanning from negative to positive potential, two peak centered at -0.54 V and -0.41 V were detected and the height of those peak increased with the growth of *Bacillus subtilis* C9. To further confirm the DPV peak assignments, riboflavin was added to the bioreactors (Fig. 2B). The -0.54 V and -0.41 V peak currents increased linearly with increasing the concentration of riboflavin (Inset curve a and b Fig. 2B), These phenomena were consistent with previous reported on *Shewanella oneidensis* MR-1 extracellular electron transfer (38). Besides, the free and cytochrome-bound riboflavin in the medium of the bioreactor with *Bacillus subtilis* C9 was investigated by the two linear relationships (R^2^ = 0.999, R^2^ = 0.999) between the peak currents at -0.54 V, -0.41 V with the adding concentration of riboflavin (Inset column a and b Fig. 2A). It was shown from inset column a Fig. 2 A that the concentration of cytochrome-bound riboflavin increased from 49.8 nM at 100 h to 192.7 nM at 172 h. the concentration of free riboflavin increased from 45.3 nM at 100 h to 306.3 nM at 172 h. These data further implied that *Bacillus subtilis* C9 might secret riboflavin.

**Fig. 2.**
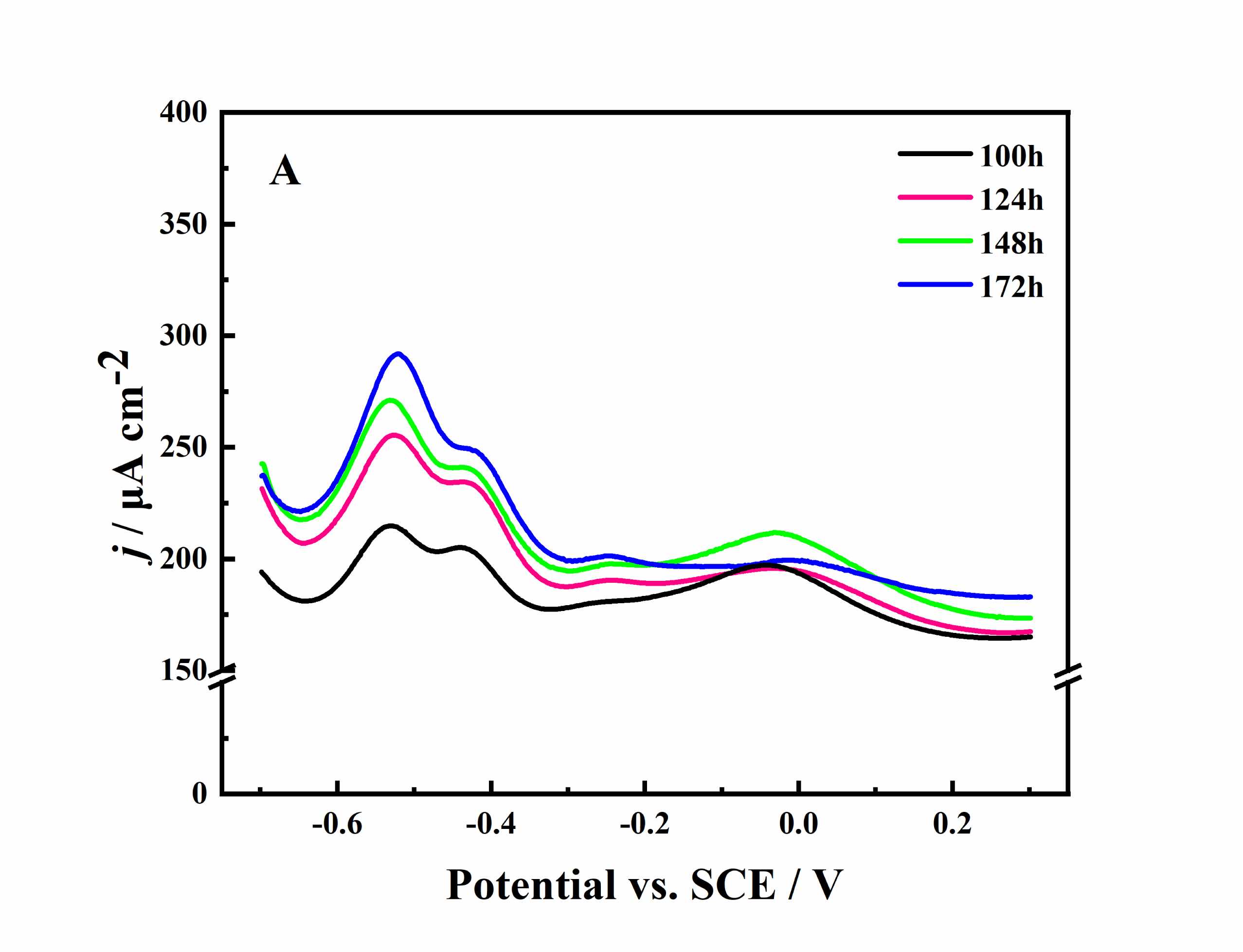
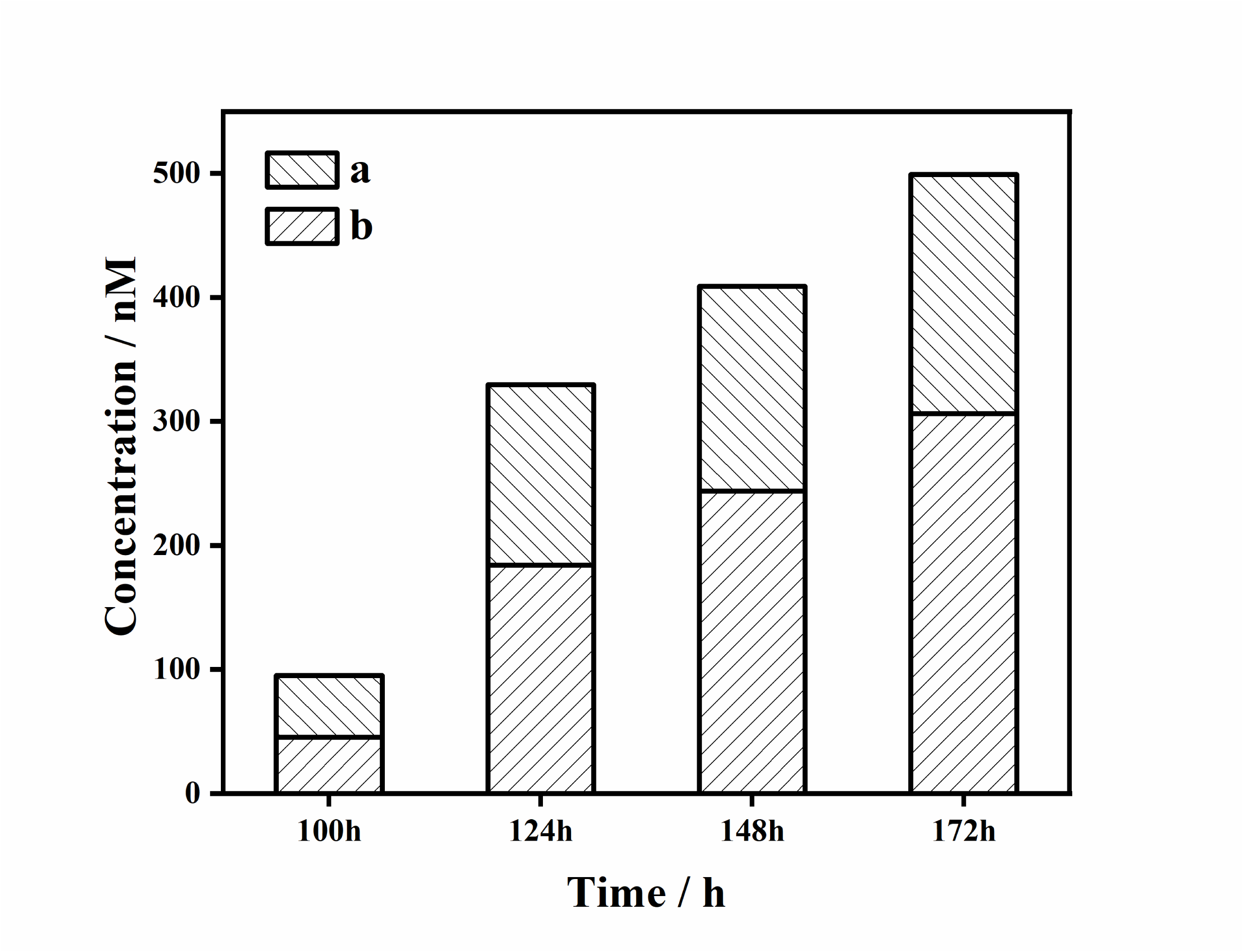
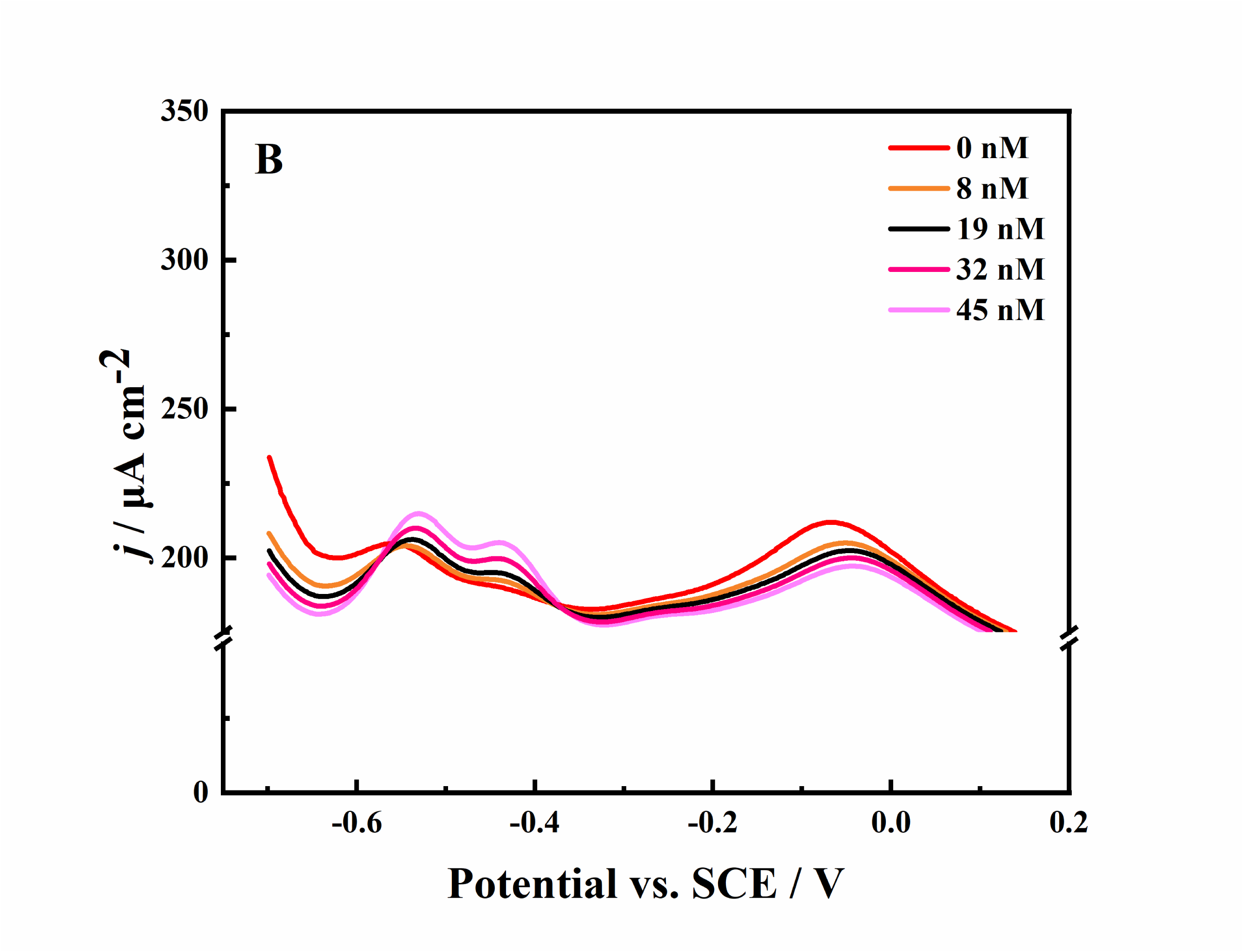
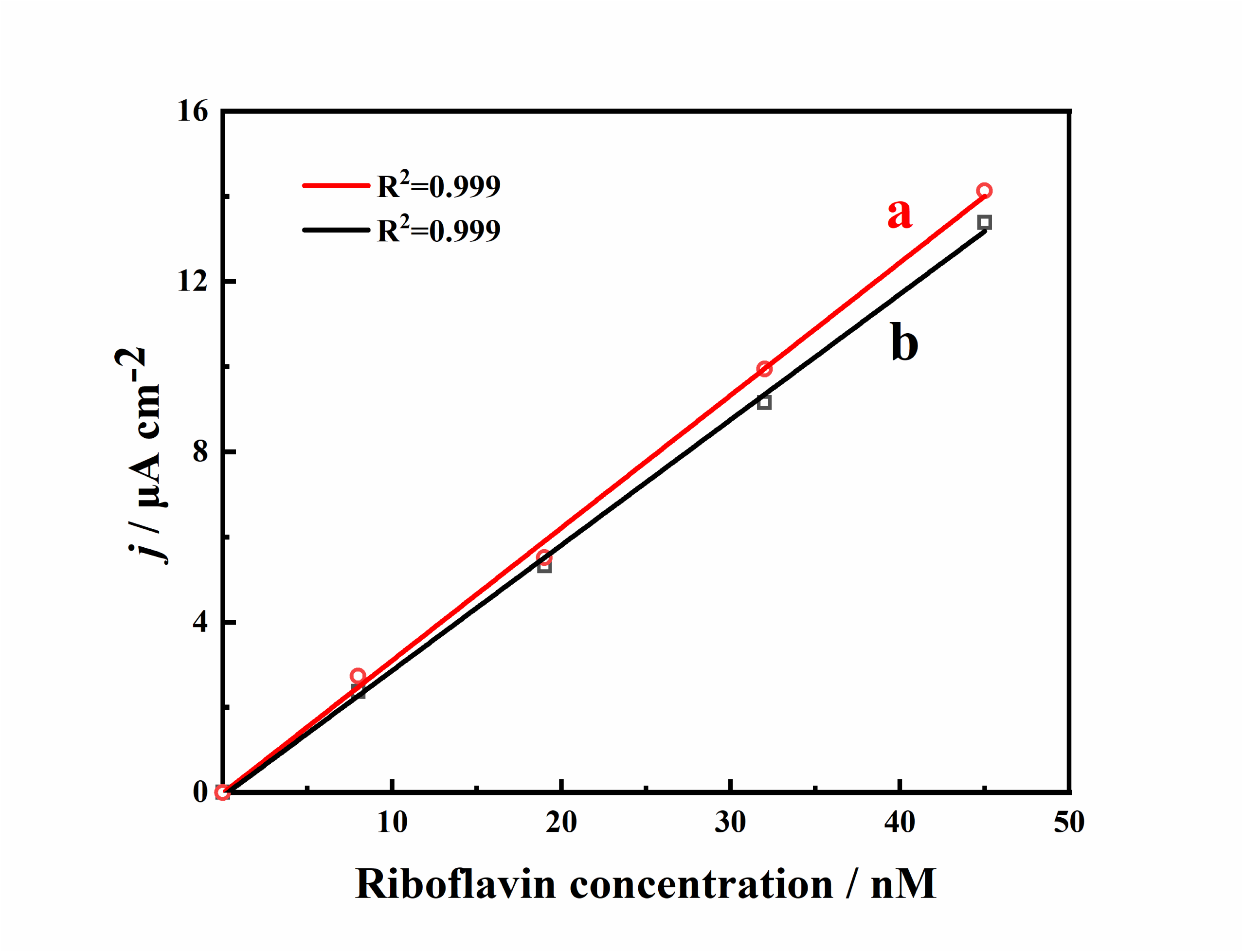
Electrochemical analysis of *Bacillus subtilis* C9. (A) DPV measured at 100,124,148 and 172 h. Inset: the dependence of free and cytochrome-bound riboflavin concentrations on time. (B) Effect of adding different concentration of riboflavin on the DPV profiles. Inset: the standard curve of the peak currents at -0.54 V(a), -0.41 V (b) versus the adding concentration of riboflavin.

### Fluorescence spectrometry analysis of strain *Bacillus subtilis* C9

The existence of riboflavin in the bioreactor with *Bacillus subtilis* C9 was proved by fluorescence spectrometry (Fig. 3 A, B and C). The fluorescence intensity of riboflavin was detected in the excitation spectra at 370 nm and 450 nm (Fig. 3 A) and emission spectra at 530 nm (Fig. 3 B and C, curves a and b). It can be seen that the data are consistent with the previous findings that *G. sulfurreducens* secreted riboflavin during growth in anaerobic medium (39). These data confirmed that the existence of riboflavin in the medium of *Bacillus subtilis* C9, which will contribute to improve electron transfer. Furthermore, it also explains the current enhancement in ternary culture (Fig. 1).

**Fig. 3.**
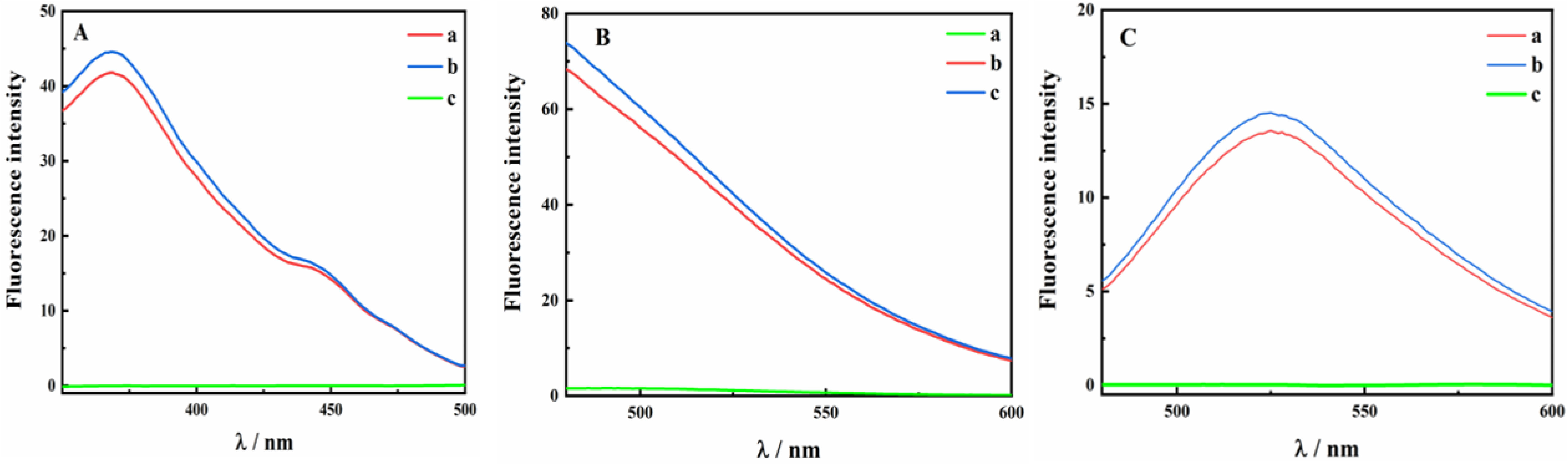
Detection of flavins secreted by *Bacillus subtilis* C9 in bioreactor. (A) Excitation spectra and (B) and (C) emission spectra of riboflavin in bioreactor, the emission wavelength at 530 nm, the excitation wavelength at 370 nm and 450 nm, the water was as the control (curve c).

### Analysis of substrate consumption and flavin secretion

It has been reported that bacterial exchange valuable and essential metabolites for the benefit of other species was key to building microbial consortia (40). The phenol-sulfuric acid method was used to analyze the cellulose degradation, while the main metabolic compositions of CMC (acetate acid) were quantified by HPLC. As shown in Fig. 4 A, the degradation of cellulose in E-BLG was higher than that in E -BG and E -LG within 120 h, which might be beneficial from the synergistic degradation of cellulose by *Cellulomonas* Lsc-8 and *Bacillus subtilis* C9. Furthermore, the concentration of acetate in these bioreactors were quantified by HPLC (Fig. 4A). Different concentration of acetate was detected in metabolites of co-cultures and ternary culture after 24 h running, which was used to produce electricity by *G. sulfurreducens* PCA. The concentration of acetate in E -BLG at 72 h (173.11±1.52 mg/L) was even higher to that in E -BG (85.91±9.97 mg/L), which was consistent with the highest current generation and biomass in the electrode biofilm of E-BLG (Fig. 1 and Fig. S6). At the end of the 120 h operation, the concentration of cellulose in E-BLG was only remained 37.10 ± 2.5 mg/L, which is lower than 43.24 ± 1.7 mg /L and 46.47 ± 1.3 mg/L in bioreactors of E -BG and E -LG. These results indicated that ternary culture could efficiently transfer essential metabolites to form a cross-feeding microbial consortium for power generation.

**Fig. 4.**
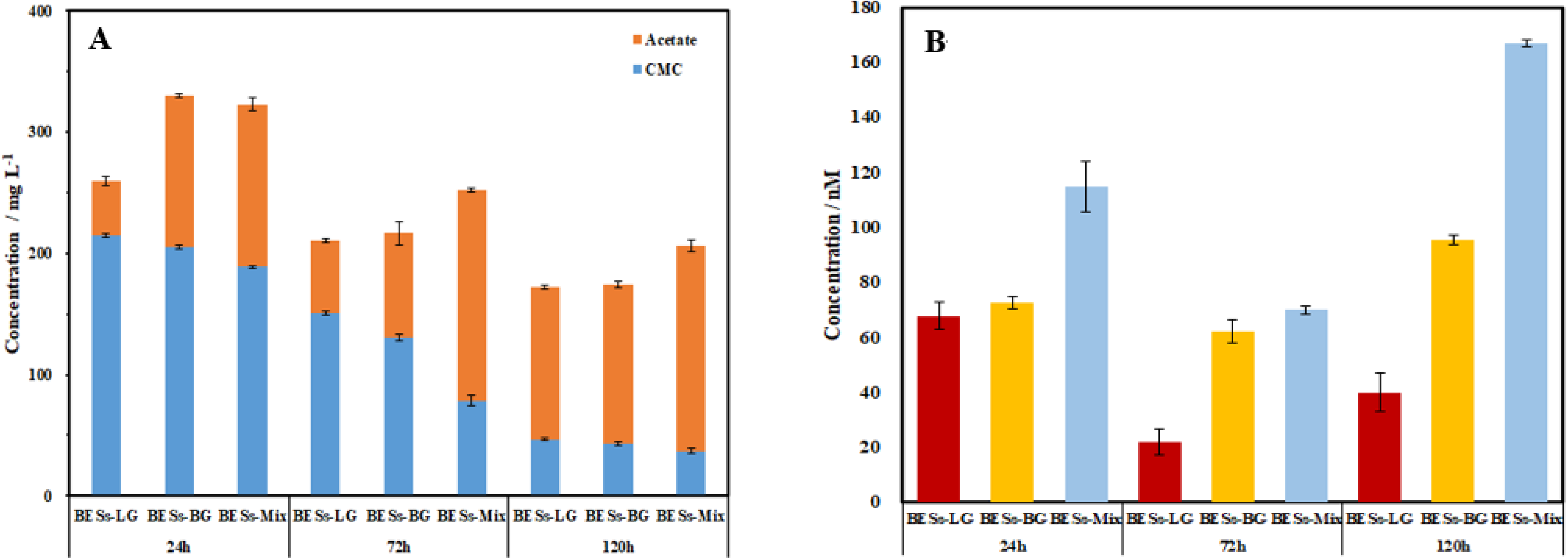
The concentration of primary metabolites and riboflavin at time 24, 72, and 120 h. (A) carboxymethyl cellulose consumption and its primary metabolites. (B) Riboflavin concentration of the bioreactors inoculated with different inoculums.

Furthermore, the concentration of riboflavin in different bioreactors were also measured by HPLC (Fig. 4 B). The concentration of riboflavin in bioreactor of E-BLG increased continuously from 115 ± 7 nM at 24 h to 167 ± 1.22 nM at 120 h, which was increased to 4.1 and 1.8 times than that in E-LG and E-BG at the same time, which further confirm *Bacillus subtilis* C9 secretes flavins (Fig. 3 and Fig. 4 B). It was reported that the extracellular electron-transfer efficiency was further promoted by microbial extracellular polymers substances (41). Therefore, riboflavin as microbial extracellular secretion to enhanced the extracellular electron transfer efficiency of EAB. It was further explaining the reason of electricity generation enhancement in ternary culture (Fig. 1). These results indicated that acetate was an essential substrate for electricity generation and riboflavin was one of the main factors improving the electricity conversion in the ternary culture.

### Community composition analysis of the defined co-cultures and ternary culture biofilm on the electrode

The high-throughput sequencing was used to detect the relative abundance of biofilm in bioreactors with different cultures (E-LG, E-BG, and E-BLG) (Fig. 5). The sample pretreatment was operated as shown Supporting Information. The abundance of *Geobacter* in biofilms with different cultures (E-LG, E-BG, and E-BLG) were 98.86%, 96.95% and 92.22%, respectively. The results showed that *Geobacter sulfurreducens* PCA was the dominant species in those biofilms. At present, It has been reported that *Geobacter* was the dominant genus in biofilms due to the competitive pressure such as fermentative and nitrate respiration (42–44). As our expectation, *Cellulomonas* Lsc-8, and *Bacillus subtilis* C9 accounted for 2.22% and 5.56% of the total abundance in the ternary culture biofilm, which were higher than their percentages in the co-culture biofilms (E-LG and E-BG), respectively. The main reason for this phenomenon might be the increased proportion of these two bacteria in biofilms improved the degradation of cellulose and produced acetic acid for the growth of *Geobacter sulfurreducens* PCA.

**Fig. 5.**
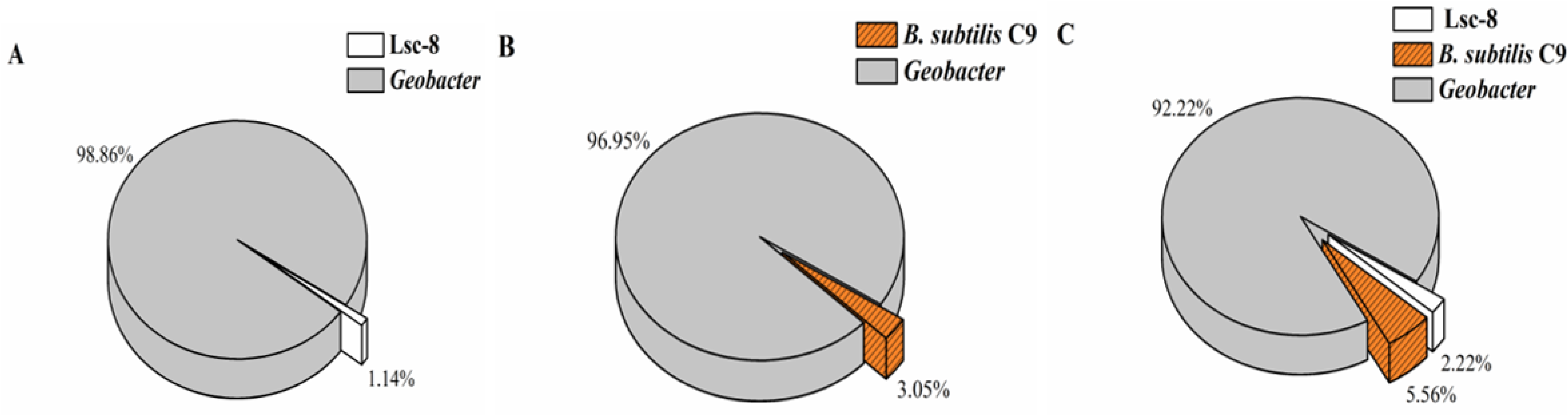
The abundant ratio of biofilm in different bioreactors (A, E-LG, B, E-BG, and C, E-BLG) (at the genus level).

### Analysis of the biomass and morphological of the defined co-cultures and ternary biofilm

The performance of bioreactors is largely associated with the bacterial loading capacity on the electrode surface. Here the biomass on electrode were quantified from different bioreactors when the highest current densities were reached (Fig. 1). The linear relationship (R^2^= 0.9925) between protein concentrations and solution absorbances at 595 nm with protein concentrations of 0 - 100 μg·mL^-1^ were shown in Supporting Information Fig. S5. It was shown from Fig. S6 that the biomass of biofilm in E-BLG was 779.46 ± 6.56 μg**·**mL^-1^, the biomass of biofilms in E-BG and E-LG were 605.14 ± 0.98 μg**·**mL^-1^ and 483.78 ± 1.65 μg**·**mL^-1^, respectively. The biomass from the biofilm in E-BL was 272.52 ± 3.53 μg**·**mL^-1^, which was much lower than that in other bioreactors. The changes in biomass of biofilms were consistent to the electricity generation performance of the defined co-cultures and ternary system. These data suggested that the ternary culture might can synergistically form dense and efficient biofilm.

The EABs on the different bioreactors was studied by Confocal scanning. The morphologcial characteristics of co-cultures and ternary biofilms were shown in Fig. 6. Confocal images of biofilm in E-BLG showed dense biofilm formation (42.68 μm) and the highest activity so that had higher current generation than E -BG and E -LG (Fig. 6 C, F and Fig. 1). Confocal images of biofilm in E-BG showed porous and the thickness of biofilm was observed 35.8 μm (Fig. 6 B and E). This might be due to cell size, production of extracellular polymeric substances or inter-species interaction (34, 45). The thickness of biofilm in E-LG was achieved 31.32 μm, which was thinner and less coverage of the electrode compared to other bioreactors (Fig. 6 A ,D and Fig. 1). These observations were very consistent with the highest biomass of ternary culture (Fig. S6) and further indicated that ternary culture can synergistically form dense and highly efficient biofilm. The results also demonstrate that *Cellulomonas* Lsc-8 and *Bacillus subtilis* C9 have positive effect on the biofilm growth of ternary culture.

**Fig. 6.**
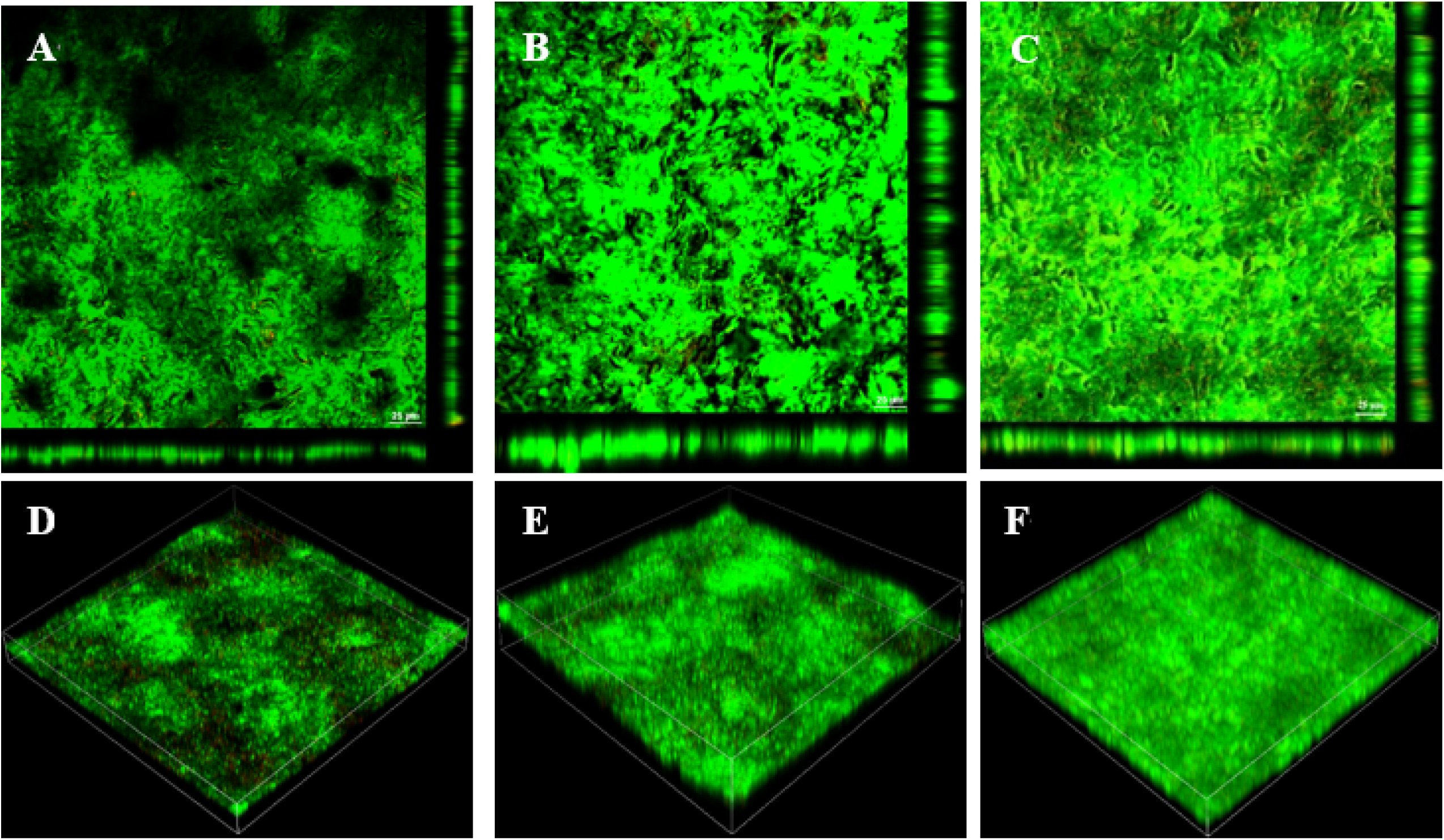
CLSM images of electrode biofilm on different bioreactors (E-LG, E-BG, and E-BLG) when achieved the highest current density, respectively. (A and D, sample from E-LG; B and E, sample from E-BG; C and F, sample from E- BLG).

### Electrochemical impedance spectroscopy analysis

Electrochemical impedance spectroscopy is a promising experimental approach in biological applications, such as the research on living cells and voltage loss (46). EIS were conducted when the bioreactors using different inocula reached the maximum current densities, the Nyquist plots were shown in Fig. S7, while the values were presented in Table S5. The working electrode charges transfer resistances (*R_ct_*) were measured as 284 Ω and 239.1 Ω for E-LG and E-BG, respectively. The bioreactor inoculated with *G. sulfurreducens* PCA using acetate was found to achieve *R_ct_* values of 105.2 Ω. The *R_ct_*value of E-LG was significantly increased to 1.19, 3.11, and 2.70 times than that of others bioreactors (E- BG, E-BLG, and E-G), respectively. It might be due to the biofilm formed by E-LG was thinner and has lower biomass (Fig. S6. and Fig. 6 A).

However, the bioreactor inoculated with ternary culture using CMC was found to achieve *R_ct_* values of 91.18Ω, which were significantly lower to that of others bioreactors. It could be attributed to the biofilm was enhanced the concentration of riboflavin and biomass in the bioreactor by the synergistic interaction between bacteria (Fig. 4 B and Fig. S7). It has been report that the synergistic interaction between bacteria enhanced the production of electron shuttles(47). A noticeably lower total resistance was observed for the E-BLG (R = 157.44 Ω) than for the E-LG (R = 324.51 Ω), E-BG (R = 268.36 Ω), and E-G (R = 173.33 Ω), respectively. There were many factors to influence the diffusion resistance, such as electrolytes, the structure of electrodes and biofilm(48). These data illustrated that the total resistance of ternary culture using CMC was lowest and further explain the current enhancement.

### 3.8. Optimization of the mixed ratio of Lsc-8, B. subtilis C9, and G.sulfurreducens PCA for electricity generation

Based on the above results, strain Lsc-8*, B. subtilis* C9*, and G.sulfurreducens* PCA can work synergistically together to generate electricity with CMC as the sole carbon source. Nine typical inoculated volume ratios were chosen to optimize the mixed ratio of ternary culture for electricity generation. The maximum electricity generation capability using CMC by ternary culture was investigated as shown in Figure S8. As proportion of *G.sulfurreducens* PCA decreased, the maximum current densities were gradual declined. The maximum current density of 796 ± 30 μA cm^2^ was obtained at the ratio 1:1:5 of strain Lsc-8, *B. subtilis* C9, and *G.sulfurreducens* PCA , which was higher than that other ratios and single *G. sulfurreducens* PCA with acetate. It can be seen that the adjustable ratio of ternary culture offers an effective method for enhancing the degradation efficiency of CMC in bioreactors. Therefore, our ternary culture system can improve effective cellulose degradation by adjusting the ratio of ternary culture composition according the properties of wastewater.

### Effect of exogenous riboflavin on electricity generation of ternary culture

To confirm whether further increasing the concentration of riboflavin can improve the electricity generation, the various concentrations of riboflavin (480, 640, 800, 960 nM and 200, 300, 400 μM) were added to the ternary culture system to investigate the impact on the bioelectricity generation (Fig. S9). At any concentration of riboflavin greater than 480 nM, there was no positive impact for the bioelectricity generation. Particularly, the initial exogenous riboflavin concentration more than 200 μM, the maximum current of ternary culture using CMC was significantly reduced. It was reported that ROS produced by photosensitized riboflavin rendered the redox status of bacterial into a compromised state leading to destroy membrane integrity ultimately causing bacterial death. In addition, photosensitized riboflavin can be used to prevent the growth of biofilms (49). Therefore, the concentration of riboflavin had important role on power generation from cellulose by ternary microbial consortium. These results also indicated that the addition of riboflavin does not further increase the electricity production of cellulose-fed ternary culture. It will be enhanced using synthetic biology approaches in the future to design and construct more robust consortia to perform complicated tasks.

This research designed and investigated a ternary culture of *Cellulomonas* Lsc-8, *Bacillus subtilis* C9 and *G. sulfurreducens* PCA as biocatalysts for cellulose-derived electricity generation. The maximum current density was 796 ± 30 µA cm^-2^, which were higher than other defined co-cultures system and that of a previously reported co-culture systems. The changes in the concentrations of CMC and acetic acid with operation time in bioreactors were also detected. *Bacillus subtilis* C9 demonstrated the ability to degrade CMC to produce acetic acid and secrete riboflavin. The highly efficiency of electricity generation in ternary system mainly benefited from the “division-of-labor” based cooperation of three species and the increase of riboflavin concentration after the introduction of *Bacillus subtilis* C9. Those results also indicated that the cellulose degradation, acetate production rate, and the concentration of riboflavin were not the main factors limiting the electricity production of cellulose-fed ternary culture following the introduction of *Bacillus subtilis* C9. This work will clarify the operation mechanisms of complicated microbial communities and select suitable microorganisms to construct more complex microbial communities for energy generation in the future.

## MATERIALS AND METHODS

### Inoculum sources and growth media

*Cellulomonas* Lsc-8 collected from rumen content as reported earlier studies in our group. *Bacillus subtilis* C9 and *Bacillus subtilis* 168 were preserved in the laboratory of the Sheep Genetic Improvement and Biological Breeding Team (30). *Cellulomonas* Lsc-8 and *Bacillus subtilis* C9 were cultured with the same growth medium. The growth medium contained (per liter) 1 g NaHCO_3_, 0.495 g KH_2_PO_4_, 0.495 g NH_4_Cl, 0.93 g NaCl, 0.066 g CaCl_2_·2H_2_O, 0.078 g MgCl_2_·6H_2_O, 0.653 g K_2_HPO_4_·3H_2_O, 5 mL of trace mineral solution, 10 mL of vitamin solution, 4 g CMC and oxygen were used as electron donor and acceptor, respectively. All media were adjusted to pH 7.2 prior to sterilization and culture growth was conducted at 30 ± 1 °C air or anaerobically. The medium of bioreactors were the same as the growth medium of *Cellulomonas* Lsc-8 without oxygen described above.

The effect of the addition of riboflavin on the electricity generation of the ternary culture was studied by adding various concentrations of riboflavin, the medium of bioreactors was the same as the medium described above. Meanwhile various initial riboflavin concentration (from 480 nM to 400 μM) medium were obtain by adding various concentrations of riboflavin (prepared by dissolving riboflavin in water).

*G. sulfurreducens* PCA was purchased from Deutsche Sammlung von Mikroorganismen und Zellkulturen GmbH. The growth medium contained (per liter) 0.6 g KCl, 1.5 g NH_4_Cl, 0.3 g KH_2_PO_4_, 0.1 g MgCl_2_**·**6H_2_O, 0.1 g CaCl_2_**·**2H_2_O, 4.64 g fumaric acid, 1.64 g sodium acetate and 10 mL of trace mineral solution, and 10 mL of vitamin solution, as the previously described(50).

For inoculum preparation, the pure cultures of bacteria were enriched in their growth medium (2% v/v) and followed by 3 days incubating with a rotary shaker at 150 rpm and 37 °C. For biofilm formation, the cell concentrations were measured by UV spectrophotometer at OD_600_ nm (Cary 60 UV-Vis). The ratio of inoculated cell densities in different combinations was shown in the Table S1. The defined various co-cultures and ternary system were named E-LB, E-LG, E-BG, and E-BLG in the following study. For the ratio optimization of ternary culture components, the ratio of Lsc-8, *B. subtilis* C9, and *G.sulfurreducens* PCA inoculation was shown in the Table S2.

### Half-cell construction and electrochemical analysis

Half-cell construction was carried out on a 16-channel potentiostat in three-electrode set-ups, including a working electrode, a counter electrode and a saturated calomel electrode (SCE, Hg/Hg_2_Cl_2_ saturated KCl, +0.244 V *vs.* hydrogen standard electrode (SHE)) as reference electrode (51). All potentials refer to SCE unless otherwise stated. The graphite plate at a dimension of 1.0 cm × 0.8 cm × 0.2 cm was used as the working electrode and current densities were normalized based on the geometric surface area (52). All electrochemical experiments were recorded using a 16-channel potentiostat in three-electrode system. The chronoamperometry measurements were anaerobically conducted in 20 mL medium under potential of 0.3 V at temperature of 30±1℃ (12). The cyclic voltammetry (CV) was measured at scan rate of 5 mV·s^-1^ from 0.3 V to -0.6 V. Differential pulse voltammetry (DPV) was carried out at scan rate of 4 mV·s^-1^ from -0.7 V to 0.3 V (19). Electrochemical impedance spectroscopy (EIS) data were used to analyze the internal resistance of bioreactors that inoculated with different inoculums, with a frequency ranging from 100 kHz∼10 MHz and the potential of - 0.45 V. All analysis were carried out for three times.

### Determination of cellulase activity

The 3,5-dinitrosalicylic acid (DNS) colorimetric method was utilized to measure cellulase activity (51). The supernatant of the growth medium was used as the crude cell-free enzyme solution after centrifugation (10,000 g, 10 min at 4 °C). A 0.5 mL aliquot of crude enzyme was added to wells containing 2.0 mL 50 mM NaAc buffer (CMC-Na, 1% (W/V)). After 15 min of incubation at 50 °C, 2.0 mL of DNS was added to each reaction and incubated at 95 °C for 5 min. Finally, each sample was diluted with water and was measured at the absorbance of 540 nm. One unit is defined as the amount of enzyme that liberates 1 µmol of reducing sugars per min, which is expressed in U/mL.

### Analysis and determination of riboflavin and metabolic products

To further analyze the ability of *Bacillus subtilis* C9 secreted riboflavin, fluorescence spectrophotometer (F-4600 series, Hitachi) was adopted to detect the supernatant solution of BES-B after operation 100 h. Riboflavin was measured by using HPLC with a UV detector and assayed by a reverse-phase C18 column (53). All samples were analyzed in triplicate.

The phenol-sulfuric acid method was used to analyze residual cellulose in each batch (54). Filtered samples (0.22 nitrocellulose filter) were analyzed for glucose and acetate acid using HPLC (Agilent 1260) with a refractive index detector and with a reverse-phase column C18 (Waters) (55).

### Community composition analysis

The biofilm samples were collected with a sterile blade when those EABs were stabilized at their maximum current density in each batch, respectively. High-throughput pyrosequencing was used to evaluate the community diversity of EABs on carbon electrodes with different co-cultures and ternary culture (E-LG, E-BG, and E-BLG). Universal primers, including the forward primer 341 F (CCCTACACGACGCTCTTCCGATCTGCCTACGGGNGGCWGCAG) and reverse primer 805 R (GACTGGAGTTCCTTGGCACCCGAGAATTCCAGACTACHVGGGTATCTAATCC), were used to amplify the V3-V4 region of 16 S *rRNA* gene. The detailed method was shown in Supporting Information.

### Determination of the biomass

The biomass of cells attached on the electrode surface was tested by the staining Coomassie brilliant blue at the point with the highest current density (19), respectively . For various EABs (E-BL, E-LG, E-BG, E-BLG), the cells attached on the working electrode surface were collected by ultrasonic treatment of 10 min. The biomass was estimated using the standard curve of protein versus absorbance at 595 nm. Each measurement was repeated 3 times.

### Morphological analysis

When the current density of the EABs reached the maximum value, the morphological analysis of EABs were studied on a Nikon A1R confocal laser scanning microscope (CLSM, Nikon, Japan) using the previous method (56). Immediately following harvesting, the planktonic cells attached on the EABs were removed by washing in 50 mM PBS for 20 min and the EABs were rinsed two times. Then those EABs were stained with the Live- dead BacLight Bacterial Viability Kit (Invitrogen) and incubated for 30 min in the dark. Those EABs were rinsed with growth medium after incubating and placed on a microscope slide. The stains were excited by two laser wavelengths, 488 and 561 nm.

## Acknowledgements

This work was financially supported by project from the National Natural Science Foundation of China (No. 21375107), the Key Science and Technology Program of Shaanxi Province, China (2023-YBNY-105).

The authors declare that they have no known competing financial interests or personal relationships that could have appeared to influence the work reported in this paper.

